# Human Pumilio proteins directly bind the CCR4-NOT deadenylase complex to regulate the transcriptome

**DOI:** 10.1101/2020.11.17.387456

**Authors:** Isioma I.I. Enwerem, Nathan D. Elrod, Chung-Te Chang, Ai Lin, Ping Ji, Jennifer A. Bohn, Yevgen Levdansky, Eric J. Wagner, Eugene Valkov, Aaron C. Goldstrohm

## Abstract

Pumilio paralogs, PUM1 and PUM2, are sequence-specific RNA-binding proteins that are essential for vertebrate development and neurological functions. PUM1&2 negatively regulate gene expression by accelerating degradation of specific mRNAs. Here, we determined the repression mechanism and impact of human PUM1&2 on the transcriptome. We identified subunits of the CCR4-NOT (CNOT) deadenylase complex required for stable interaction with PUM1&2 and to elicit CNOT-dependent repression. Isoform-level RNA sequencing revealed broad co-regulation of target mRNAs through the PUM-CNOT repression mechanism.

Functional dissection of the domains of PUM1&2 identified a conserved N-terminal region that confers the predominant repressive activity via direct interaction with CNOT. In addition, we show that the mRNA decapping enzyme, DCP2, has an important role in repression by PUM1&2 N-terminal regions. Our results support a molecular model of repression by human PUM1&2 via direct recruitment of CNOT deadenylation machinery in a decapping-dependent mRNA decay pathway.

## Introduction

RNA-binding proteins (RBPs) are crucial for regulating gene expression (Gerstberger, Hafner, and Tuschl 2014; Gehring, Wahle, and Fischer 2017). Human PUM1 and PUM2 (PUM1&2) are members of the conserved Pumilio and Fem-3 binding factor (PUF) family of RBPs that typically repress gene expression (Wickens et al. 2002; Goldstrohm, Hall, and McKenney 2018). PUM1&2 exhibit marked specificity for the consensus RNA sequence 5′-UGUANAUA, the Pumilio Response Element (PRE) (Zamore, Williamson, and Lehmann 1997; X. Wang et al. 2002). They regulate PRE-containing mRNAs from diverse genes, predominantly acting as repressors that cause degradation of target mRNAs and reduce protein production (Morris, Mukherjee, and Keene 2008; Van Etten et al. 2012; Bohn et al. 2018; Goldstrohm, Hall, and McKenney 2018; Wolfe et al. 2020; Yamada et al. 2020).

PUM1&2 have essential regulatory functions (Wickens et al. 2002; Arvola et al. 2017; Goldstrohm, Hall, and McKenney 2018) and simultaneous inactivation of both is embryonically lethal in mice (Zhang et al. 2017; Lin et al. 2018; Uyhazi et al. 2020). PUM1&2 control growth, hematopoiesis, neurogenesis, behavior, fertility, and neurological functions (Goldstrohm, Hall, and McKenney 2018; Lin et al. 2019; Uyhazi et al. 2020). They have been implicated in cancer (Kedde et al. 2010; Miles et al. 2012; S. Lee et al. 2016; Bohn et al. 2018; Naudin et al. 2017; Yamada et al. 2020), neurodegeneration, and epilepsy (Gennarino et al. 2015; Follwaczny et al. 2017; Gennarino et al. 2018). Recently, mutations in PUM1 were linked to the disorders PUM1-associated developmental disability, ataxia, and seizure (PADDAS) and PUM1-related adult onset, cerebellar ataxia (PRCA) (Gennarino et al. 2015, 2018).

Given that all vertebrates have two Pumilio paralogs (Goldstrohm, Hall, and McKenney 2018), do PUM1&2 have redundant, overlapping, or unique regulatory functions? *In vitro* RNA-binding studies showed that PUM1&2 display highly correlated specificities (Cheong and Hall 2006; Lu and Hall 2011; Van Etten et al. 2012; Jarmoskaite et al. 2019; Wolfe et al. 2020). RNA co-immunoprecipitation analysis demonstrated significant overlap of bound mRNAs but also provided evidence for unique subsets (Morris, Mukherjee, and Keene 2008; Galgano et al. 2008; Hafner et al. 2010; Yamada et al. 2020). Phenotypic analysis of knockout mice lends support for both overlapping and unique functions (Goldstrohm, Hall, and McKenney 2018; Lin et al. 2018, 2019; Uyhazi et al. 2020). While distinct expression patterns are likely relevant, both PUM1&2 are coincidentally expressed in a broad array of tissues and cell types (Goldstrohm, Hall, and McKenney 2018; Spassov and Jurecic 2002). Systematic analysis of the individual and combined functional impact of human PUM1&2 on the transcriptome is necessary to address this important consideration.

Pumilio proteins from species ranging from insects to mammals share a primary structure that includes a large N-terminal extension and a C-terminal RNA-binding domain (RBD) (Goldstrohm, Hall, and McKenney 2018). The RBD was structurally characterized and its RNA-binding affinity and specificity have been intensively studied (Lu, Dolgner, and Hall 2009; X. Wang et al. 2002). The RBD contributes to repression by antagonizing the Poly-Adenosine Binding Protein (PABP) and also by recruiting RNA decay factors (Kadyrova et al. 2007; Goldstrohm and Wickens 2008; Van Etten et al. 2012; Weidmann et al. 2014). However, in fruit flies, the RBD of Pumilio makes a minor contribution to repression (Weidmann and Goldstrohm 2012; Weidmann et al. 2014), which prompted us to examine the structure and function of human PUMs.

The N-terminal regions of PUM1&2 are less conserved among orthologs and show no similarity to other proteins, but do carry 70% identity between PUM1&2 paralogs (Weidmann and Goldstrohm 2012; Goldstrohm, Hall, and McKenney 2018). In *Drosophila* Pumilio, three N-terminal repression domains (RD1, 2 and 3) were identified (Weidmann and Goldstrohm 2012). In that same study, the N-terminal regions of human PUM1&2 were shown to have repressive activity when artificially directed to an mRNA in *Drosophila* cells. However, the function of the PUM1&2 N-terminal regions in translational repression and mRNA degradation remained untested in human cells.

The CCR4-NOT (CNOT) complex catalyzes the shortening of the 3′ poly(A) tail of mRNAs in a process termed deadenylation, and plays a pivotal role in initiating translational repression and mRNA decay (Goldstrohm and Wickens 2008). Importantly, the repressive activity of PUM1&2 was reduced when deadenylation was inhibited (Van Etten et al. 2012), suggesting a functional connection between PUMs and CNOT; however, the broader impact of this on the human transcriptome remained unknown.

The eight subunit human CNOT complex contains two distinct deadenylase enzymes, Pop2-type paralogs CNOT7 or CNOT8, and Ccr4-type paralogs, CNOT6 or CNOT6L (Goldstrohm and Wickens 2008). CNOT1 serves as a scaffold for additional CNOT subunits (i.e. CNOT2, 3, 9, 10, and 11) that mediate protein interactions with RBPs, microRNA-induced silencing complex, mRNA decay factors, and translational regulators (Fabian et al. 2013; Chen et al. 2014; Bhandari et al. 2014; Sgromo et al. 2017; Mathys et al. 2014; Raisch et al. 2018, 2019). Multiple CNOT components were reported to copurify with human PUMs, including the deadenylases (Goldstrohm et al. 2006; Van Etten et al. 2012), but precise contacts were not mapped and functional roles of the CNOT components in PUM-mediated repression remained unknown.

In this study, we interrogate the role of the CNOT complex in repression by human PUM1&2, and find that several subunits are necessary for repressive activity. We then perform transcriptome-wide, isoform-level RNA sequencing and discover hundreds of target mRNAs that are coregulated by PUM1&2 and CNOT. We map the major repressive domains of PUM1&2, identify their direct interactions with the CNOT complex, and show that the mRNA decapping pathway plays an important role in the PUM-CNOT repression mechanism.

## Results

### Compositional CNOT requirements for PUM-mediated repression

We and others have shown that human PUMs can accelerate degradation of PRE-containing mRNAs (Morris, Mukherjee, and Keene 2008; Bohn et al. 2018; Goldstrohm, Hall, and McKenney 2018; Wolfe et al. 2020) and implicated the CNOT deadenylase complex in this mechanism (Van Etten et al. 2012). However, the functional requirement for CNOT and the molecular basis of the PUM1&2-mediated repression remained unknown. To dissect the role of each CNOT subunit in PUM repression, we utilized PRE-containing Nano-luciferase (Nluc) reporter genes (**Figures 1A,B**) (Van Etten et al. 2012; Bohn et al. 2018). Cotransfected Firefly luciferase (Fluc) was used to normalize for variation in transfection efficiency. Repressive activity was measured by calculating the fold change of the Nluc 3xPRE reporter relative to a version wherein the 5′-UGU trinucleotide of the PRE that is essential for PUM-binding was substituted by 5′-ACA (Nluc 3xPRE mt) (Van Etten et al. 2012; Bohn et al. 2018). As expected, depletion of PUM1&2 (**Figure 1C**) alleviated PRE-mediated repression in human HCT116 cells, whereas the non-targeting control (NTC) siRNAs had no effect (**Figure 1D**).

**Figure 1.**
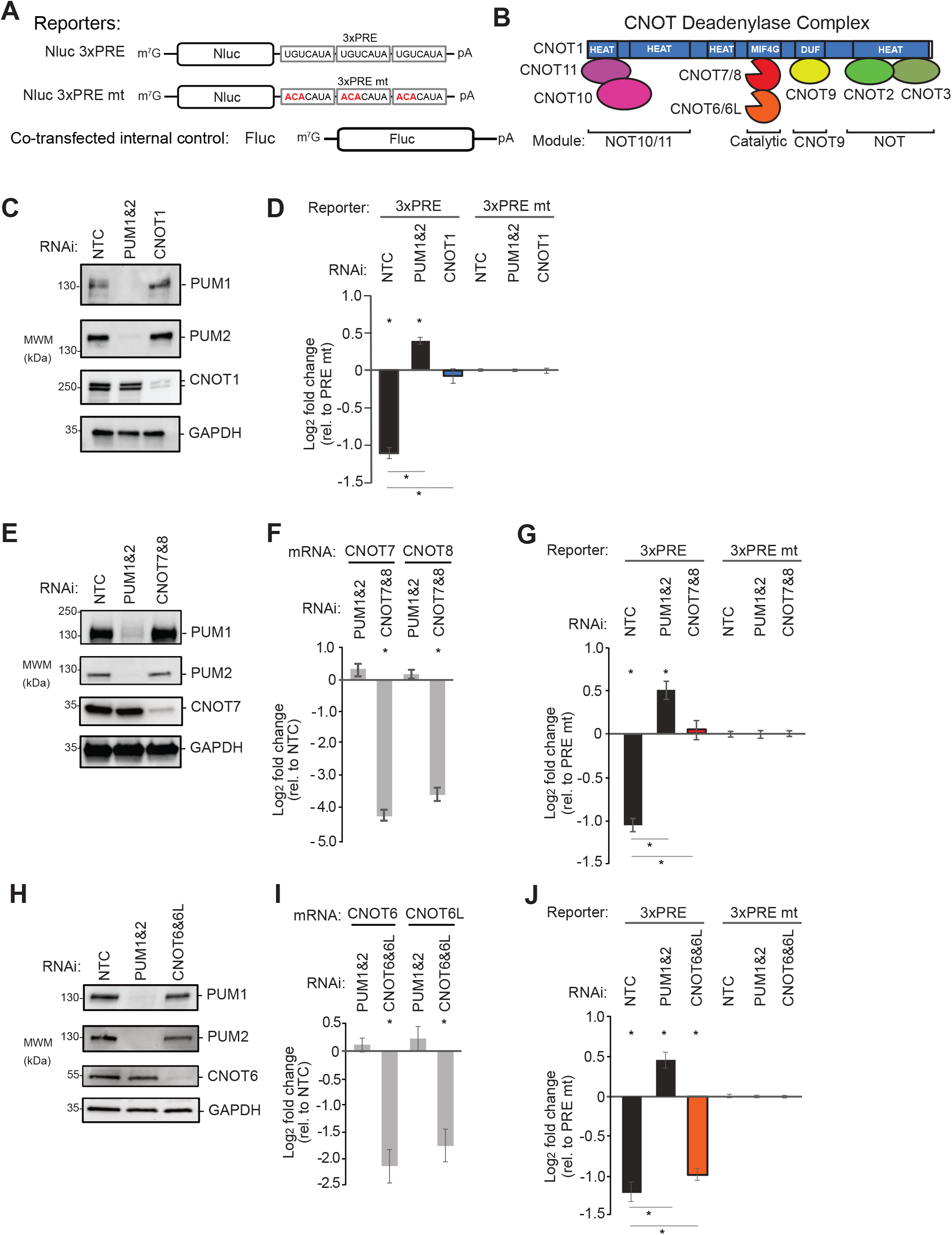
Compositional CNOT requirements for PUM-mediated repression. Plotted data, listed in **Table S1**, are graphed as mean log2 fold change values +/− SEM. **A**. Luciferase reporter constructs with Nluc followed by a 3′UTR containing 3 copies of WT or mutant (mt) Pumilio Response Element, PRE. Fluc served as control. **B**. Architecture and module organisation of the human CNOT complex. **C**. Western blot of PUM1&2 and CNOT1 confirming RNAi depletion in HCT116 cells. GAPDH served as loading control. NTC: non-targeting control siRNA. **D**. Effect of depletion of PUM1&2 or CNOT1 on repression of the Nluc 3xPRE, relative to the Nluc 3xPRE mt reporter. NTC RNAi serves as a control. The ‘*’ symbols denote a p-value < 0.05 relative to the negative control MS2-HT (above x-axis) or between the indicated RNAi conditions (below the x-axis). N=9. **E**. Western blot of RNAi depletion of CNOT7 and PUM1&2. **F**. RT-qPCR confirmed RNAi depletion of CNOT7&8 mRNAs, relative to NTC. The ‘*’ symbols denote a p-value of less than 0.05. N=9. **G**. Effect of RNAi depletion of CNOT7&8 and PUM1&2 on PRE-mediated repression. N=9. **H**. Western blot of RNAi depletion of CNOT6 and PUM1&2. **I**. RT-qPCR measurement of RNAi-mediated depletion of CNOT6&6L mRNAs. N=9. **J**. Effect of RNAi-mediated depletion of CNOT6&6L and PUM1&2 on PRE-mediated repression.

We first depleted the central scaffold CNOT1 protein (**Figure 1C**) and observed a complete loss of PRE-PUM repressive activity (**Figure 1D**). Next, we systematically depleted the deadenylase enzymatic subunits. Efficient depletion was verified by western blot for CNOT7 (**Figure 1E**) as well as RT-qPCR for both CNOT7 and CNOT8 (**Figure 1F**). Knockdown of CNOT7 and CNOT8 alleviated PRE-mediated repression (**Figure 1G)**. Depletion of both CNOT7 and CNOT8 was necessary due to their functional redundancy (Lau et al. 2009; Yi et al. 2018). Knockdown of the two Ccr4-type deadenylases, CNOT6 and CNOT6L (**Figures 1H,I**), resulted in a modest but statistically significant reduction in repressive activity (**Figure 1J**). Depletion of CNOT2 slightly reduced PRE-PUM repressive activity (**Figure S1A-B**). In contrast, depletion of CNOT3 (**Figure S1A-B**), CNOT9, CNOT10, or CNOT11 (**Figure S1C-F**) did not significantly impact repressive activity. These results reveal that the deadenylase enzyme subunits and central scaffold of the CNOT complex are necessary for PRE-PUM mediated repression in human cells.

### The PUM-CNOT axis regulates a substantial proportion of the human transcriptome

We then set out to determine the impact of the PUM-CNOT repression mechanism on the transcriptome. First, we identified the mRNAs regulated by each PUM independently and in combination, revealing that they coregulate many RNAs. We utilized Poly(A)-ClickSeq (PAC-Seq), an RNA sequencing approach that measures differential mRNA expression of 3′UTR isoform variants (Routh et al. 2017; Elrod et al. 2019), to analyze RNA isolated from cells depleted of PUM1, PUM2, or both. Efficient depletion of PUM1&2 was verified by western blot and was also observed in the PAC-Seq data (**Figure 2A,B**). To assess significant differences in gene expression, a 1.3-fold change cutoff in mRNA level (calculated for each RNAi condition relative to negative control RNAi condition, NTC) was used with an adjusted p-value < 0.05 (**Table S2**).

**Figure 2.**
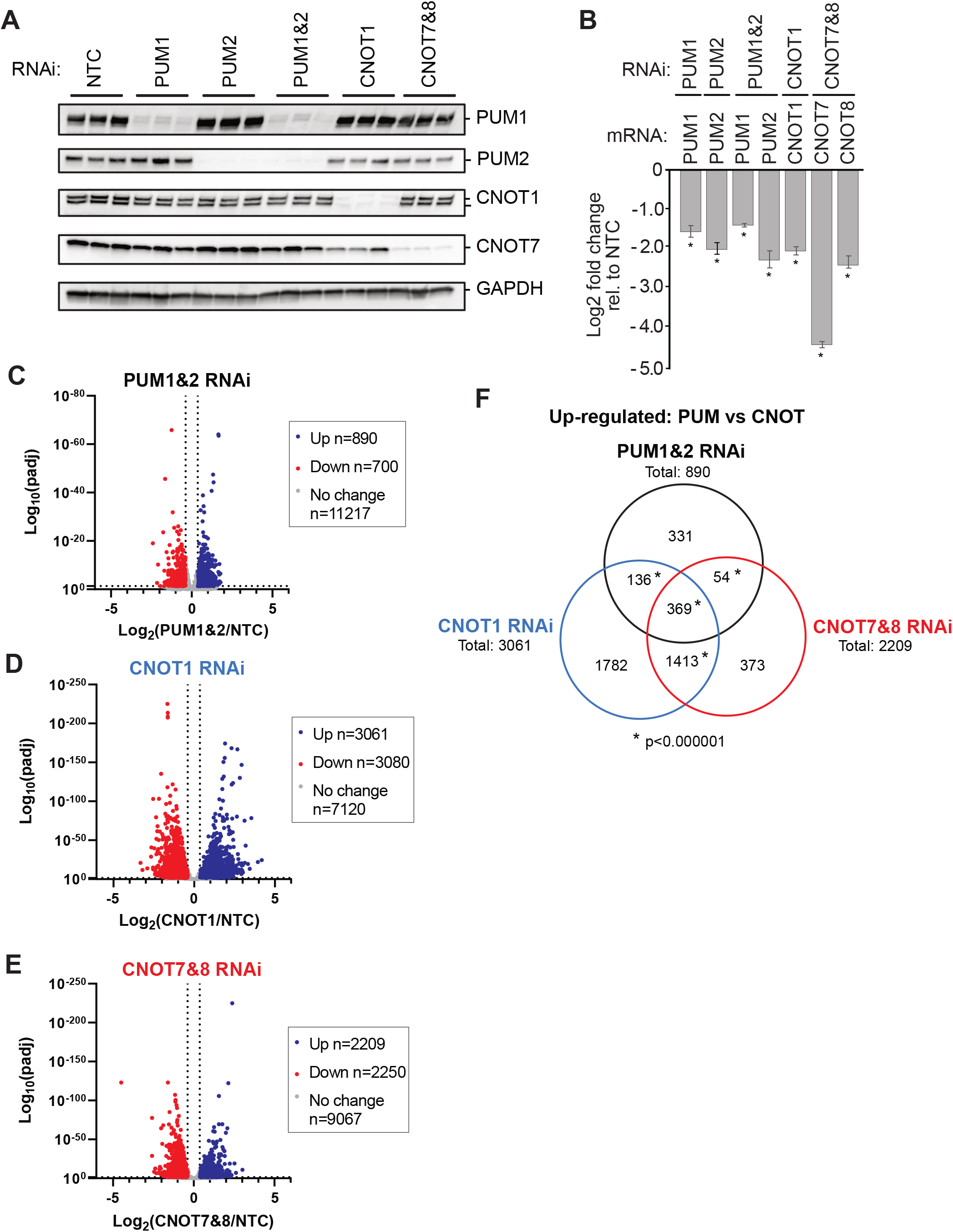
The PUM-CNOT axis regulates a substantial proportion of human transcriptome. **A**. Western blot of RNAi depletion of PUM1, PUM2, or both simultaneously (PUM1&2), CNOT1, or CNOT7 and CNOT8 (CNOT7&8), from three biological replicates of HCT116 cells used for PAC-Seq analysis. Cells treated with non-targeting control (NTC) siRNAs served as a negative control. GAPDH served as loading control. **B**. RNAi depletion of PUM1, PUM2, CNOT1, CNOT7, and CNOT8 mRNAs measured in the PAC-seq data. Mean log2 fold change values +/− SEM (**Table S1**) are plotted relative to NTC. The ‘*’ symbols denote a p-value < 0.05 relative to the NTC control. N=3. **Panels C-E.** Volcano plots of statistical significance (adjusted p-value) versus mean log2 fold change of RNA levels, relative to NTC, measured by PAC-seq for each RNAi condition: **C.** PUM1&2 RNAi; **D.** CNOT1 RNAi; **E.** CNOT7&8 RNAi. The number of genes that were up- or down-regulated by 1.3-fold or greater with an adjusted p-value < 0.05 are reported in the inset box. PAC-Seq data and statistics are reported in **Table S2**. **F.** Venn diagram of gene sets that were significantly up-regulated in each indicated RNAi condition. The ‘*’ symbol for each comparison denotes a p-value of < 0.000001 (hypergeometric test for multi-set overlap). Gene sets and statistics are reported In **Table S3**.

Simultaneous depletion of PUM1&2 altered the expression of 1590 genes, and we focused on the 890 genes that were upregulated in a manner consistent with PUM-mediated repression (**Figure 2C**). Depletion of either PUM1 or PUM2 individually, however, upregulated 132 or 102 genes, respectively (**Figure S2A-C**). Comparison of the gene sets that are upregulated by PUM1, PUM2, and PUM1&2 RNAi indicates that the majority of PUM-repressed genes are detected only when both are depleted (721 genes)**(Figure S2D, Table S3)**. Most of the remaining genes in the PUM1&2 gene set overlap with the individual PUM1 and/or PUM2 knockdowns (105 or 146, respectively), and a small subset were exclusively upregulated by either PUM1 (27 genes) or PUM2 (20 genes).

To identify direct targets of PUM1&2-mediated repression, we compared the upregulated gene set (**Figure S2E: “PUM1&2 RNAi”**) with previously identified genes that contain PREs (**Figure S2E: “PRE”**) and that are bound by PUMs (**Figure S2E: “Bound”**)(Galgano et al. 2008; Morris, Mukherjee, and Keene 2008; Hafner et al. 2010; Bohn et al. 2018). The majority of the PUM1&2 repressed genes have a consensus PRE (521 genes) and/or were found to be bound by PUMs (383 genes). When these three stringent criteria were collectively applied, we identified 335 direct PUM targets (**Figure S2E, Table S3**).

Emerging evidence indicates that PUM1&2 can alternatively promote expression of certain genes (Naudin et al. 2017; Bohn et al. 2018; Wolfe et al. 2020), referred to as PUM-mediated activation (Goldstrohm, Hall, and McKenney 2018). Here, we identified 62 genes that were downregulated by depletion of PUM1&2, contain PREs, and are bound by PUMs (**Figure S2F, Table S3**), providing additional support for direct PUM-mediated activation. This data will facilitate future efforts to elucidate the mechanism (Bohn et al. 2018; Goldstrohm, Hall, and McKenney 2018).

Based on their crucial roles in PUM-mediated repression, we evaluated the effects of depletion of CNOT1 and CNOT7&8 on mRNA levels in the same PAC-Seq experiment. As anticipated for factors that have broad roles in mRNA decay, their depletion (**Figure 2A, B**) upregulated the levels of many genes including upregulation of 3061 for depletion of CNOT1 and 2209 for depletion of CNOT7&8 (**Figure 2D,E**). To identify PUM1&2 repressed target genes that are also repressed by CNOT, we compared the upregulated gene sets from PUM1&2, CNOT1, and CNOT7&8 RNAi conditions. The PUM1&2 repressed genes overlap with 505 genes that are upregulated by CNOT1 depletion and 423 genes by CNOT7&8 depletion (**Figure 2F**). Further comparison revealed that 369 genes are upregulated in all three RNAi conditions. Of those, 215 contain a consensus PRE, 161 are bound by PUMs, and 138 genes met all three criteria (**Figure S2G, Table S3**). Together, these results identify many endogenous mRNAs that are repressed by PUM1&2 and the CNOT complex, providing new insight into the global impact of the PUM-CNOT repression mechanism.

### The RNA-binding domains of PUM1&2 are necessary but not sufficient for repression

To understand how PUM1&2 cause CNOT-mediated repression, we dissected their repressive domains. First, we tested the contributions of their RBD and N-terminal regions (**Figure 3A**). We isolated this analysis to exogenously provided PUM1&2 by utilizing an altered specificity reporter gene assay wherein the RNA recognition motif of the sixth repeat of each PUM was altered to specifically bind a mutated PRE that contains a 5′-UGG motif in place of the wild type 5′-UGU (**Figure 3B**, Nluc 3xPRE UGG), as previously established (Van Etten et al. 2012; Weidmann and Goldstrohm 2012; Weidmann et al. 2014).

**Figure 3.**
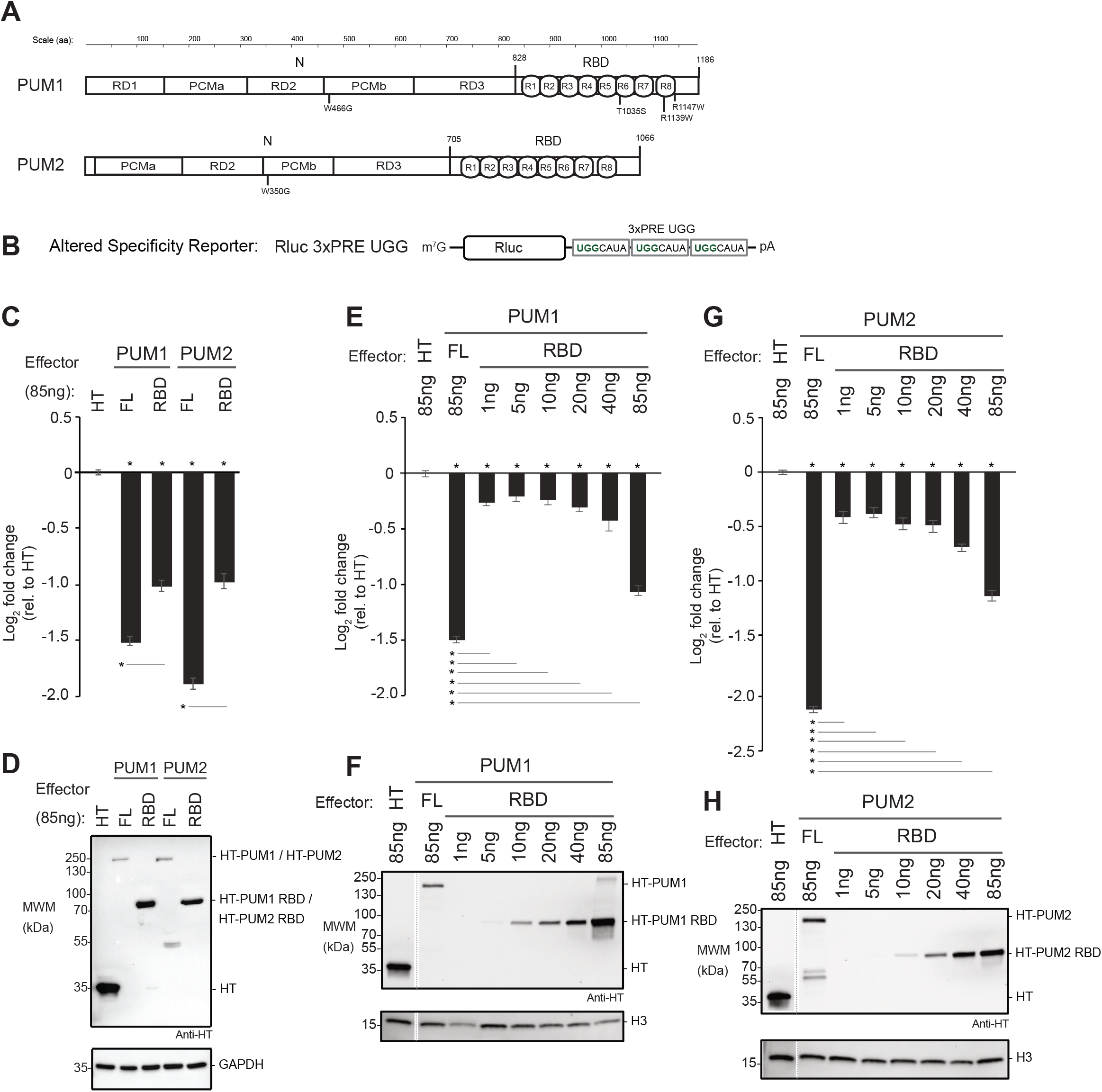
The RNA-binding domains of PUM1&2 are necessary but not sufficient for repression. Mean log2 fold change values +/− SEM are plotted and listed in **Table S1**. The ‘*’ symbols denote a p-value < 0.05 relative to the negative control MS2-HT (above x-axis) or between the indicated proteins (below the x-axis). **A**. Domain organization of PUM1 and PUM2 indicating the N-terminal region (N) with putative Repression Domains (RD), Pumilio Conserved Motifs (PCMa and PCMb), and RNA-Binding Domain (RBD) with 8 PUF repeats (R1-8) (Weidmann and Goldstrohm 2012; Goldstrohm, Hall, and McKenney 2018). Residue numbering on top. Putative cap binding residues, W466 and W350, are indicated below (Cao, Padmanabhan, and Richter 2010). The PUM1 residues linked to the diseases PADDAS, R1139W and R1147W, and PRCA, T1035S, are indicated below. **B**. Altered specificity assay reporter gene, Nluc 3xPRE UGG, that specifically measures the activity of exogenous PUM1&2 (R6as) engineered to bind to the UGG motifs. **C**. Altered specificity reporter assay comparing the repressive activity of full length (FL) or RBD PUM1 or PUM2 R6as effectors, relative to negative control HT. N=9. **D**. Western blot of HT-tagged proteins from a representative experimental replicate from samples from panel C. GAPDH served as loading control. **E**. Comparison of RBD to full length (FL) PUM1 (R6as) using the altered specificity reporter assay. Total mass of transfected DNA was balanced across transfections with an empty vector. N=9. **F**. Western blot of HT-tagged PUM1 proteins in panel E from a representative experimental replicate. Histone H3 served as loading control. **G**. Same as Panel E, except using PUM2 FL and RBD (R6as) proteins. N=9. **H**. Western blot of HT-tagged PUM2 proteins used in panel G from a representative experimental replicate.

Full length PUM1&2 both had significantly greater repression activity than their corresponding RBDs (**Figure 3C**). Notably, the RBD constructs were more abundantly expressed than the full length PUM1&2 (**Figure 3D**). In light of this observation, we compared the repressive activity and expression level across a titrated range of transfected RBD plasmid relative to full length PUM1 (**Figure 3E,F**) or PUM2 (**Figure 3G,H**). This titration allowed a more accurate assessment that, when expressed at near equal level to the corresponding full length protein, each RBD was a much weaker repressor. This analysis confirms that, while the RBD is necessary, it is not sufficient for full PUM1&2 function, suggesting that the N-terminal regions have important repressive activity.

### N-terminal regions of human PUMs are potent repressors

We then focused on the potential activity of the PUM1&2 N terminal regions. Little information about these regions was available, with the exception of a small conserved motif previously reported to be involved in translational repression by the *Xenopus* PUM2 ortholog (Cao, Padmanabhan, and Richter 2010). That motif was shown to bind the 7-methyl guanosine cap. We therefore tested the potential role of this motif by substituting the critical cap-binding tryptophan by glycine (**Figure 3A**, PUM1 W466G, PUM2 W350G) and measuring the effect on PUM1&2 activity using the altered specificity reporter assay. Neither W466G nor W350G substitution in PUM1 and PUM2, respectively, alleviated repression (**Figure S3**). Thus, the cap-binding motif does not contribute to repressive activity.

To dissect the repressive activity of the N-terminal regions of PUM1&2, independent of their RBDs, we employed a tethered function reporter assay (Jeff Coller and Wickens 2007), wherein four binding sites for the MS2 phage coat protein are embedded in the 3′UTR of Nluc (**Figure 4A**, Nluc 4xMS2) (Abshire et al. 2018). The MS2-HT negative control did not significantly affect the Nluc 4xMS2 reporter relative to a control reporter that lacked MS2 binding sites, Nluc ΔMS2 (**Figure 4B**). In contrast, the positive control, MS2-CNOT7 deadenylase, robustly repressed Nluc 4xMS2 but not Nluc ΔMS2 (**Figure 4B**), showing that recruitment of CNOT7 is sufficient to repress expression. When the N-terminal region of either PUM (MS2-PUM1 N and MS2-PUM2 N) was expressed (**Figure 4C**), the Nluc 4xMS2 reporter protein was significantly reduced, dependent on the MS2 binding sites (**Figure 4B**). Using northern blotting, we observed that repression by both MS2-PUM1 N and MS2-PUM2 N reduced Nluc 4xMS2 mRNA levels relative to the negative control (**Figure 4D,E**). These data show that the N-terminal regions of both PUM1&2 function to repress protein and mRNA expression.

**Figure 4.**
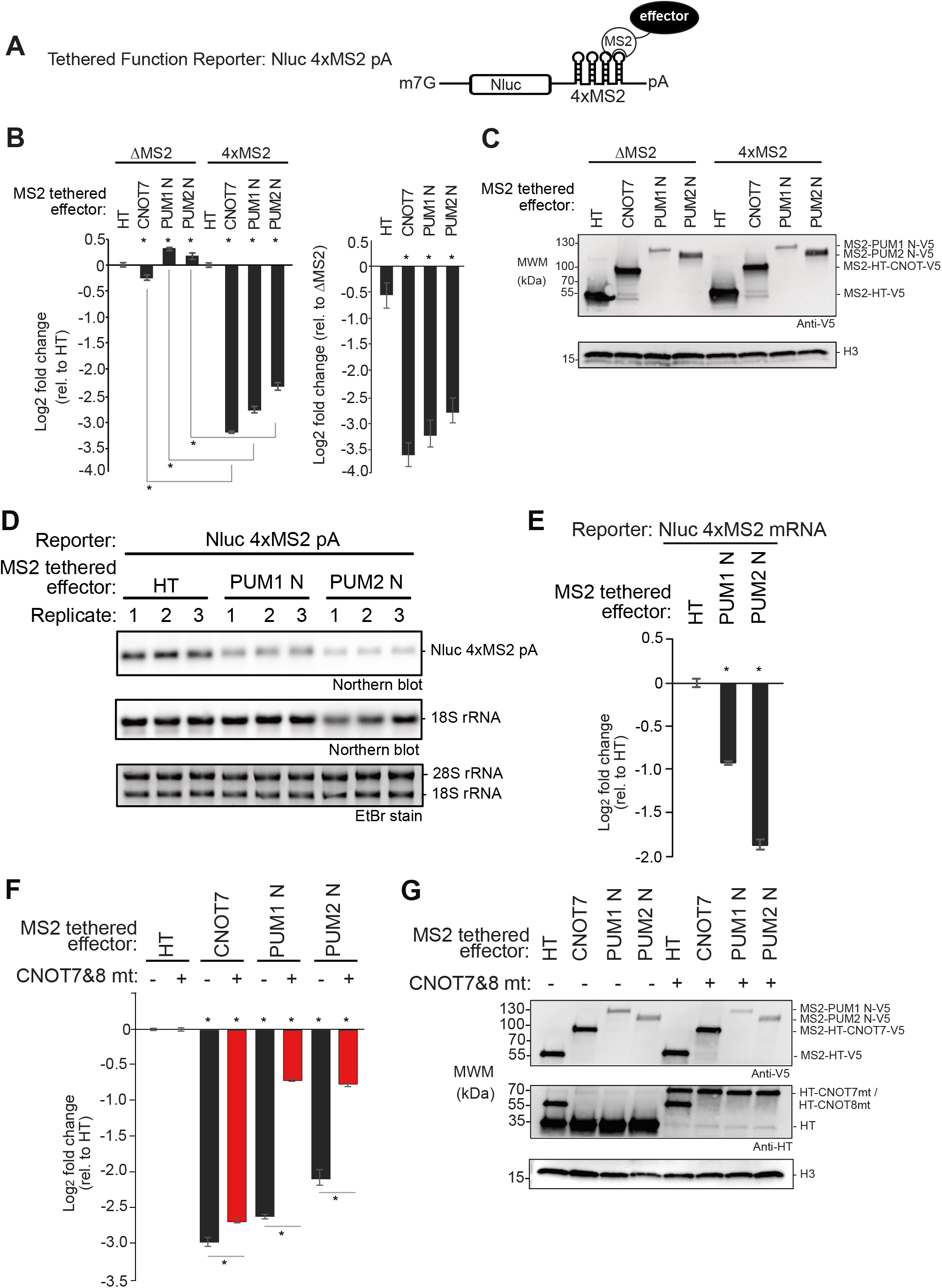
N-terminal regions of human PUMs are potent repressors in isolation. Mean log2 fold change values +/− SEM are plotted and listed in **Table S1**. The ‘*’ symbols denote a p-value < 0.05 relative to the negative control MS2-HT (above x-axis) or between the indicated conditions (below the x-axis). **A**. Tethered function reporter gene, Nluc 4xMS2 pA, containing four MS2 coat protein stem-loop binding sites in its 3′UTR and terminating in a 3′ poly(A) tail. **B**. Repressive activity of the N-terminal regions of PUM1&PUM2 (PUM1 N, PUM2 N) measured with the tethered function assay. PUM1&2 proteins were expressed as fusions to the RNA-binding domain of MS2 phage coat protein with a V5 epitope tag. MS2 fused to Halotag (HT) or CNOT7 served as negative and positive controls, respectively. Nluc ΔMS2 pA reporter, which lacked the MS2 binding sites, served as a negative control reporter gene. The left graph shows the activity of each effector relative to MS2-HT for each reporter, Nluc ΔMS2 pA or Nluc 4xMS2 pA. The right graph shows the activity of each effector on the Nluc 4xMS2 pA relative to the Nluc ΔMS2 pA. N=9. **C**. Western blot of proteins from a representative experimental replicate from samples in panel B. Histone H3 served as a loading control. **D**. Northern blot of Nluc 4xMS2 pA mRNA measured the effect of tethered MS2 fusions of PUM1 N, PUM2 N, or HT. Ethidium Bromide (EtBr) and northern blot detection of rRNA assessed RNA integrity and loading. N=3. **E**. Quantitation of the northern blot in panel D. **F**. The role of CNOT in repression by tethered MS2 fusions of PUM1 N and PUM2 N was assessed using the Nluc 4xMS2 pA reporter. CNOT activity was inhibited by overexpressing HT fusions of dominant negative mutant CNOT7&8 proteins (CNOT7&8 mt, +), compared to HT (−). Repression relative to MS2-HT negative control. N=9. **G**. Western blot of V5-tagged proteins and HT-tagged CNOT7&8 mt from a representative experimental replicate from samples in panel F.

### The PUM1&2 N-terminal regions function in repression via the CNOT deadenylase

Given that CNOT is necessary for PUM-mediated repression, we next asked if the N-terminal regions of PUM1&2 require CNOT function. We first attempted RNAi knockdown but, interestingly, CNOT depletion concomitantly decreased the expression of the transfected PUM1&2 constructs (data not shown). As an alternative strategy to block CNOT-catalyzed mRNA decay that has been used successfully (Yamashita et al. 2005; Piao et al. 2010; Temme et al. 2010; Van Etten et al. 2012; Loh, Jonas, and Izaurralde 2013; Mishima and Tomari 2016), we coexpressed a combination of catalytically inactive CNOT7&8 dominant negative mutants (or HT negative control) and then measured the effect on repression by MS2-PUM1-N and MS2-PUM2-N (**Figure 4F**). Importantly, the expression levels of PUM1&2 constructs were not altered (**Figure 4G**). We observed that overexpression of CNOT7&8 mutants elicited a pronounced, significant decrease in repression by both MS2-PUM1 N and MS2-PUM2 N relative to control (**Figure 4F**). In contrast, tethered MS2-CNOT7 was minimally perturbed by the CNOT7&8 mutants, which is an expected outcome where an active deadenylase is directly tethered to the reporter mRNA. These observations indicate that the N-terminal regions of PUM1&2 require a functional CNOT complex to repress mRNAs.

### Functional dissection of the N-terminal repressive regions of PUM1&2

Having established that the N-terminal regions of PUM1&2 have crucial repressive activity, we next mapped their repressive domains. Previously, we identified three repressive domains in *Drosophila* Pumilio (Weidmann and Goldstrohm 2012). Using sequence conservation as a guide, we delineated corresponding regions in human PUM1&2 (**Figure 3A**, RD1-3) (Weidmann and Goldstrohm 2012; Goldstrohm, Hall, and McKenney 2018). Notably, human PUM2 lacks a region corresponding to the *Drosophila* RD1.

We compared the ability of the individual RDs and RBDs of PUM1&2 to regulate the Nluc 4xMS2 reporter. We found that RD3 regions alone exhibited robust repression activity, matching that observed with the complete N-terminal region (**Figure 5A**), and exceeding that of the RBD. RD1 of PUM1 caused a moderate decrease in reporter expression whereas RD2 of PUM1 or PUM2 exhibited minimal effects. As PUM1 RD3 was expressed at a higher level than the other fragments (**Figure 5B**), we then titrated the PUM1 RD3 plasmid whilst keeping the total mass of transfected plasmid DNA constant across conditions. By this approach, we found that tethered PUM1 RD3 repression activity (**Figure 5C**) and expression level (**Figure 5D**) increased proportionally with the mass of transfected plasmid, plateauing at more than 8-fold observed level of repression. By western blot, the level of PUM1 RD3 protein at 20 ng of transfected plasmid was comparable to that of PUM1 N, RD1, or RD2 at 85 ng (**Figure 5D**). We also measured the effect of the N-terminal regions and RD3 of PUM1&2 on levels of the Nluc 4xMS2 reporter mRNA by performing RT-qPCR. All four constructs significantly reduced the reporter mRNA level relative to the negative control MS2-HT (**Figure 5E,F**). Based on these results, we conclude that RD3 of PUM1&2 confers the major repressive activity, and can function autonomously to reduce protein and mRNA levels.

**Figure 5.**
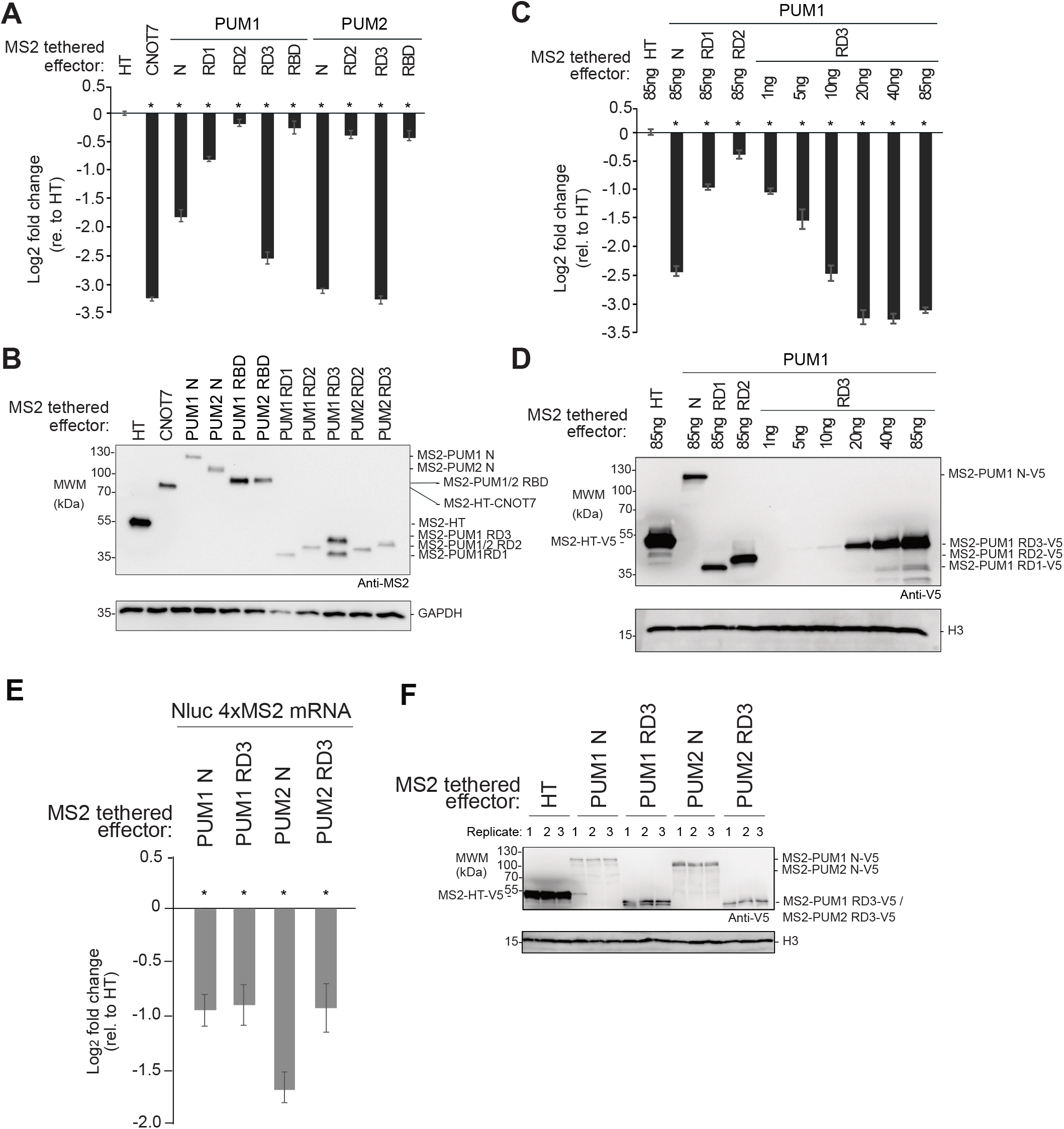
Functional dissection of the N-terminal repressive regions of human PUM1&2. Mean log2 fold change values +/− SEM are plotted and listed in **Table S1**. The ‘*’ symbols denote a p-value < 0.05 relative to the negative control MS2-HT (above x-axis) or between the indicated conditions (below the x-axis). **A**. Activity of putative repression domains (RD1, RD2, and RD3) and RBDs of PUM1&2, fused to MS2 coat protein and HT, measured in tethered function assays. MS2 fusions of HT and CNOT7 served as negative control and positive control, respectively. N=9. **B**. Western blot of MS2-tagged proteins from a representative experimental replicate from samples in panel A. GAPDH served as loading control. **C**. Activity of MS2-PUM1 RD3 compared to PUM1 N, PUM1 RD1 and PUM1 RD2 MS2 fusions relative to MS2-HT. Transfected plasmid mass is indicated and is balanced across conditions with an empty vector. N=9. **D**. Western blot of V5-epitope tagged proteins from a representative experimental replicate from samples in panel C. Histone H3 service as a loading control. **E**. Effect of PUM1 N, PUM1 RD3, PUM2 N and PUM2 RD3 MS2 fusion effectors on Nluc 4xMS2 mRNA levels, relative to MS2-HT, measured by RT-qPCR. N=9. **F**. Western blot of V5-tagged proteins from a representative experimental replicate from samples in panel E.

### Repression domain 3 of PUM1&2 binds directly to the CNOT complex

We next asked if RD3 of each PUM interacts with CNOT in cells. To test this, V5 epitope-tagged PUM1 RD3 or PUM2 RD3 ‘bait’ proteins, or V5-tagged CNOT8 positive control, were expressed in HCT116 cells and immunopurified. To disrupt potential RNA-mediated interactions, RNA in each sample was degraded by addition of RNases, as verified by gel electrophoresis (**Figure S4**). Following extensive washing, the bound proteins were eluted and probed in western blots to detect CNOT subunits. As a negative control, immunoprecipitations were performed from cells transfected with an empty expression vector (**Figure 6A**, control). We observed that multiple CNOT subunits (CNOT1, 2, 3, 6, and 7) coimmunoprecipitated with PUM1 RD3 and PUM2 RD3 and the positive control bait, but not the negative control (**Figure 6A**). Thus, RD3 of PUM1&2 associates with the CNOT complex.

**Figure 6.**
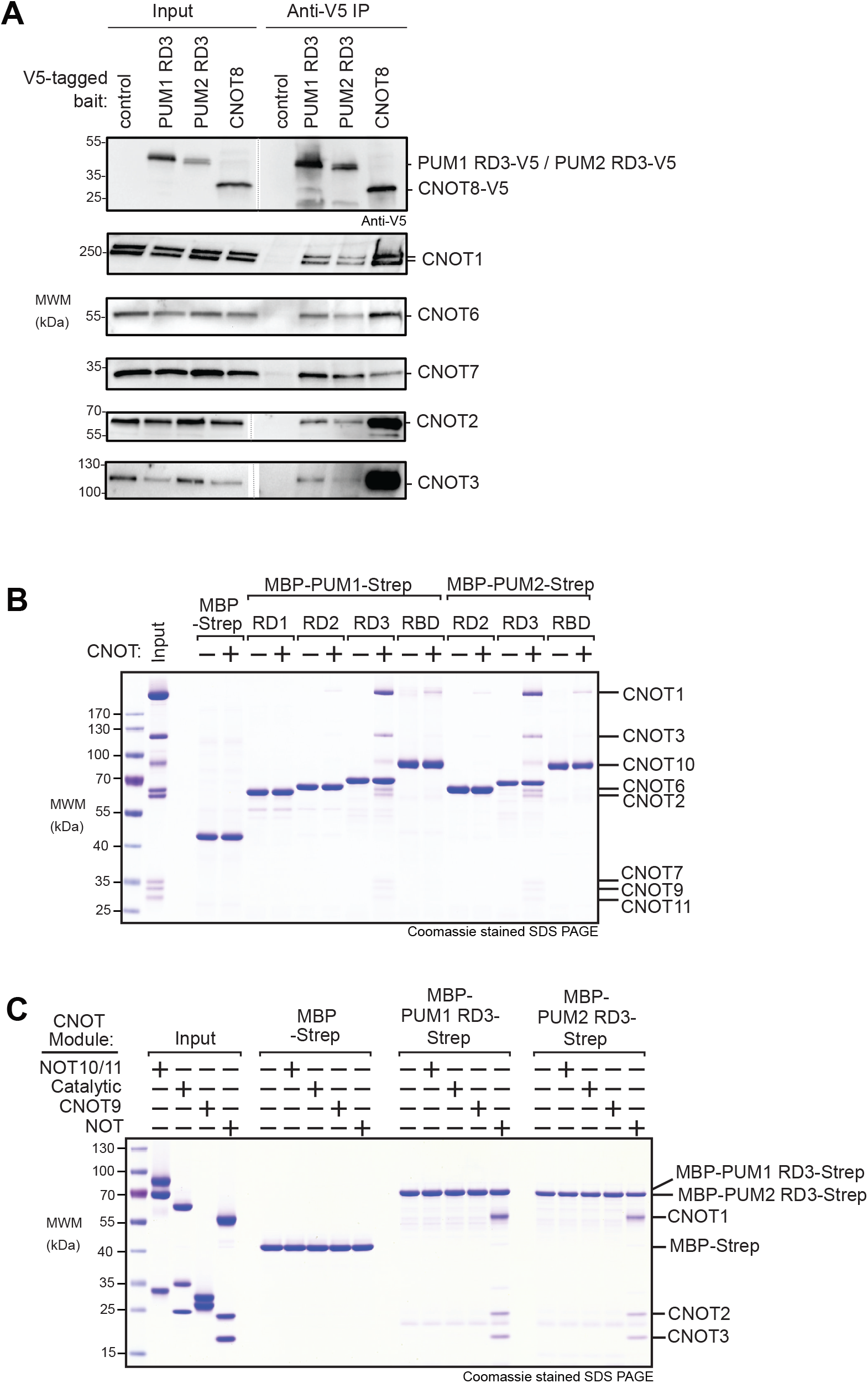
Repression domain 3 of PUM1&2 binds directly to the CNOT complex. **A**. Co-immunoprecipitation of RD3 domains of PUM1&2 proteins from HCT116 cells. CNOT8 served as a positive control and empty vector served as a negative control. Western blot of each V5-tagged bait protein in cell extracts (Input) and anti-V5 immunoprecipitates (IP). Endogenous CNOT subunits were detected by western blot. Cell extracts were treated with RNases A and 1, and RNA digestion was confirmed in Figure S4. **B**. Coomassie stained SDS-PAGE of in vitro pulldown assays with recombinant purified RD1, RD2, RD3 and RBD domains of PUM1 and PUM2, produced as fusions with Maltose Binding Protein (MBP) and StrepII (Strep) affinity tags with the intact purified CNOT complex (Input). MBP-Strep served as negative control. **C**. Same as panel C, but with the four indicated CNOT modules (Input).

We then confirmed the PUM-CNOT interaction using an independent approach and further investigated the protein-protein contacts. First, we reconstituted a recombinant CNOT complex containing all eight core subunits (Raisch et al. 2019). We then performed in vitro ‘pull-down’ assays using recombinant bead-bound PUM1&2 domains (RD1, RD2, RD3, and RBD) that had C-terminal maltose binding protein (MBP) and N-terminal StrepII (Strep) affinity tags (**Figure 6B**). Following incubation and washing, the bound complexes were eluted and analyzed by Coomassie-stained SDS-PAGE. We observed that the CNOT complex interacted with RD3 of PUM1&2, with all CNOT subunits detected (**Figure 6B**), whereas RD1 and RD2 did not appreciably interact. We also note that CNOT subunits were present in the pull-downs of PUM1&2 RBDs, albeit at a lower level.

To delineate the component(s) of the CNOT complex that directly interacts with RD3, we tested the ability of PUM1&2 RD3 regions to bind individual recombinant, purified modules of the CNOT complex using the in vitro pull-down assay. This revealed that the NOT module (Boland et al. 2013), consisting of the C-terminal CNOT1 fragment, and structured regions of CNOT2 and CNOT3, was directly bound by RD3 of both PUMs but not the MBP-Strep negative control (**Figure 6C**). In contrast, no interaction was detected with the NOT10/11, catalytic (CNOT6/7), or CNOT9 modules. This analysis precisely delineates the contacts between RD3 and the NOT module, providing a key insight into the molecular mechanism by which PUM1&2 promotes CNOT-mediated repression.

### The 3′ poly(A) tail and 3′ end accessibility are important for repression by PUM1&2 N-terminal regions

Previously we observed that the poly(A) tail was necessary for efficient repression by full length PUM1&2 (Van Etten et al. 2012). Here, we compared the ability of the N-terminal regions of PUM1&2 to repress the Nluc 4xMS2 reporter with a poly(A) tail (**Figure 4A**) to a similar reporter that terminates with a MALAT1 triple-helical RNA structure (**Figure 7A**) (Abshire et al. 2018). If deadenylation is essential for PUM-repression, then the poly(A) tail should be required. In contrast, the MALAT1 3′ end is processed to yield a triple helix structure that stabilizes an mRNA from 3′ decay whilst supporting translation (J. E. Wilusz et al. 2012; Brown et al. 2014; J. E. Wilusz 2016). Thus, if PUM1&2 retain the ability to repress the MALAT1 reporter, this would indicate an additional deadenylation-independent repression mechanism.

**Figure 7.**
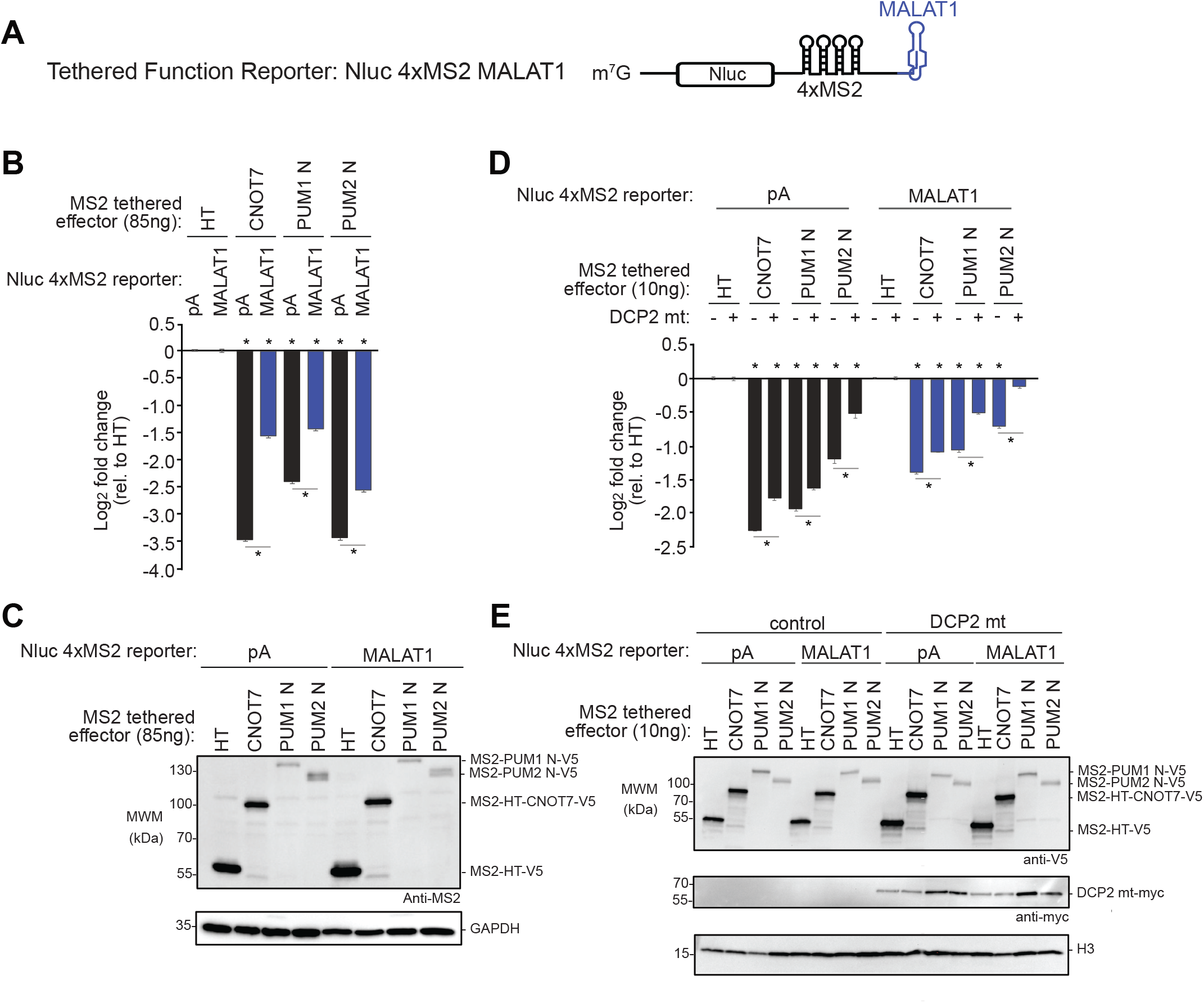
PUM1&2-mediated repression is directed via the decapping-dependent pathway. Mean log2 fold change values +/− SEM are plotted and listed in **Table S1**. The ‘*’ symbols denote a p-value < 0.05 relative to the negative control MS2-HT (above x-axis) or between the indicated conditions (below the x-axis). **A**. The Nluc 4xMS2 MALAT1 reporter gene used in the tethered function assays. The 3′ end of this reporter terminates with the MALAT1 triple helix structure. **B**. Repressive activity of MS2-fused PUM1 N, PUM2 N, or CNOT7 effector proteins measured by the tethered function assay with either Nluc 4xMS2 pA (black bars) or MALAT1 (grey) reporters, relative to MS2-HT negative control. N=9. **C**. Western blot of MS2-tagged protein from a representative experimental replicate for panel B. GAPDH served as a loading control. **D**. The tethered function assay was used to determine the effect of overexpressed, dominant negative DCP2 mutant (DCP2 mt) on repression activity of MS2-tethered PUM1 N, PUM2 N, or CNOT7 effector proteins, relative to MS2-HT negative control. Repression of Nluc 4xMS2 pA and Nluc 4xMS2 MALAT1 reporters was measured without (−) or with (+) overexpressed DCP2 mt. N=9. **E**. Western blot of V5-tagged proteins and myc-tagged DCP2 mt from a representative experimental replicate for panel D.

When tethered, the N-terminal regions of PUM1&2 exhibited significantly reduced repressive activity on the MALAT1 reporter relative to the polyadenylated reporter (**Figure 7B and 7C**), indicating that the poly(A) tail and/or the accessibility of the 3′ end are important. Consistently, repression by tethered CNOT7 was also reduced on MALAT1 versus poly(A). These results for PUM1&2 N-termini and CNOT7 effector proteins are consistent with the involvement of deadenylation of the 3′ end of the mRNA in their repressive mechanisms. The observed residual inhibitory activity on the MALAT1 reporter may be due to the ability of the CNOT complex to cause translational repression and 5′ decapping (see Discussion) (Behm-Ansmant et al. 2006; Cooke, Prigge, and Wickens 2010; Waghray et al. 2015).

### PUM1&2-mediated repression is directed via the decapping-dependent pathway

Following poly(A) tail shortening by the CNOT complex, the mRNA may be degraded by either the 5′-to-3′ decapping-dependent decay pathway or the 3′-to-5′ decay pathway (Garneau, Wilusz, and Wilusz 2007). To investigate if decapping is necessary for PUM-mediated repression, we inhibited decapping by overexpressing a dominant negative DCP2 mutant in cells (Covarrubias et al. 2011; Chang et al. 2014; Erickson et al. 2015; Sgromo et al. 2017). We then tested the ability of tethered PUM1 or PUM2 N-termini to repress the pA and MALAT1 reporters in this mutant background, compared to control. We observed that overexpression of the DCP2 mutant significantly reduced repression by each effector on both reporters (**Figure 7D,E**). These results indicate that PUM1&2-mediated repression is directed via the decapping-dependent pathway, even in the absence of the poly(A) tail and deadenylation.

## Discussion

Pumilio proteins have emerged as archetypal sequence-specific RNA-binding factors (Wickens et al. 2002; Arvola et al. 2017; Goldstrohm, Hall, and McKenney 2018). Here, we reveal new insights into PUM1&2-mediated regulation with individual mRNAs and on the global transcriptome. We mapped the domains of PUM1&2 that elicit repression, determined the complement of the requisite co-repressors, identified new domains of PUM1&2 that directly bind to the CNOT deadenylase complex, and measured the impact of the PUM-CNOT repression mechanism on the transcriptome.

### Model of PUM1&2-mediated repression

We found that the N-terminal region of PUM1&2, in particular the RD3 region, is principally responsible for PUM1&2 repressive activity, requires functional CNOT deadenylase, and can operate autonomously when directed to an mRNA. The RD3 regions of PUM1&2 are likely intrinsically disordered (Goldstrohm, Hall, and McKenney 2018) and directly bind to the C-terminal NOT1-2-3 module of CNOT. We envision that the RBD of each PUM provides RNA-binding affinity and specificity, whereas the RD3 module binds to CNOT, thus rationalizing the modular architecture of PUM1&2. Together, these interactions effectively tether CNOT directly to target mRNAs where it can shorten the 3′ poly(A) tail and initiate the degradation pathway.

Our results contribute to the growing body of evidence that the CNOT complex, and the NOT and CNOT9 modules in particular, serve as hubs for post-transcriptional regulation by sequence-specific RNA-binding regulatory factors (Goldstrohm and Wickens 2008; Temme et al. 2010) such as Roquin (Sgromo et al. 2017), TTP, microRNA induced silencing complex (Jonas and Izaurralde 2015; Fabian et al. 2013), and NANOS (Bhandari et al. 2014; Raisch et al. 2019). Interestingly, the *Drosophila* ortholog of NANOS synergizes with Pumilio to repress certain mRNAs (Weidmann et al. 2016; Arvola et al. 2017), hinting at general conservation of multivalent interactions between CNOT and specific RBPs.

As part of this study, we examined the effect of PUM1 missense mutations (**Figure 3A**, T1035S, R1139W, or R1147W) that are associated with the neurodevelopmental disorders PADDAS and PRCA (Gennarino et al. 2018). Utilizing the altered specificity approach, we observed a minor but statistically significant decrease in their repression activities (**Figure S5**). Whilst the R1147W and T1035S mutations were proposed to reduce PUM1 stability (Gennarino et al. 2018), it is possible the modest observed reduction in repression activity may be sufficient to trigger disease progression. The potential for these mutations to alter neuron-specific protein interactions or intracellular localization also warrants future study.

### Functional specificity of deadenylases in PUM1&2-mediated repression

The CNOT complex contains two types of deadenylases enzymes, Pop2-type CNOT7&8 and Ccr4-type CNOT6&6L, that contribute to general mRNA deadenylation (Goldstrohm and Wickens 2008; Yi et al. 2018; Raisch et al. 2019). Surprisingly, we observed that depletion of CNOT7&8 alleviated repression whereas the depletion of CNOT6&6L had a minor effect, indicating an apparent functional specificity for Pop2-type deadenylases in PUM1&2-mediated repression. The fact that the two deadenylase types exhibit differences in catalytic properties *in vitro (Raisch et al. 2019)* and *in vivo* (Yi et al. 2018) is potentially relevant. Further, PABP negatively affects CNOT7&8 activity whereas it promotes deadenylation by CNOT6&6L (Webster et al. 2018; Yi et al. 2018). One hypothesis that should be tested in the future is that the functional deadenylase specificity of PUM1&2 may derive from their ability to antagonize PABP (Weidmann et al. 2014).

### A role for decapping in PUM1&2-mediated repression

The observation that PUM1&2 can repress mRNAs that lack a poly(A) tail (e.g. MALAT1 or Histone Stem Loop (Van Etten et al. 2012)) suggests that there exists an additional facet to the repression mechanism. The fact that recruitment of the CNOT complex can elicit both deadenylation and deadenylation-independent translational repression is likely relevant (Cooke, Prigge, and Wickens 2010; Ozgur et al. 2015; Waghray et al. 2015; Kamenska et al. 2016; Räsch et al. 2020). Moreover, our data indicate that decapping contributes to repression by PUM1&2. In the 5′-to-3′ mRNA decay pathway, decapping follows deadenylation (Garneau, Wilusz, and Wilusz 2007); although, decapping mechanisms by RBPs have been shown to bypass deadenylation (Badis et al. 2004; Muhlrad and Parker 2005). CNOT also directly binds to decapping factors (Jonas and Izaurralde 2013; Valkov, Jonas, and Weichenrieder 2017) and the 5′ exoribonuclease XRN1, which degrades decapped mRNAs (Chang et al. 2019). By exploiting the CNOT-mediated interactions in the decay network, PUM1&2 could consequently facilitate decapping of mRNA targets, which warrants future study.

### Impact of PUM-CNOT mechanism on the transcriptome

PAC-Seq analysis shows that human PUM1&2 have a profound, wide impact on gene expression. We performed gene ontology (GO) analysis with the 890 genes that were upregulated in response to PUM1&2 depletion. The most significantly enriched GO terms include positive regulation of RNA polymerase II transcription, MAPK signaling, regulation of cell migration, endocytosis, Wnt signaling, and post-embryonic development, as well as cancer terms (**Table S3**). Many of these are direct targets of PUM1&2, based on the presence of PRE sites, evidence of binding by PUMs, and supported by comparative analyses (Bohn et al. 2018; Goldstrohm, Hall, and McKenney 2018; Wolfe et al. 2020). Combining these PAC-Seq results from HCT116 cells with data from HEK293 cells (Bohn et al. 2018; Wolfe et al. 2020) increases the number of identified PUM1&2-repressed mRNAs to 1476. Further, 369 genes are mutually upregulated by depletion of PUM1&2, CNOT1, and CNOT7&8, including known PUM targets (e.g. cell cycle factors CCNE2, CCNG2, CKS2, and cancer gene KRAS) (Bohn et al. 2018; Wolfe et al. 2020), which were enriched with similar gene ontologies as for PUM1&2 (**Table S3**). Future research should explore the implications of PUM1&2 regulation of cancer genes and pathways, expanding upon existing links (Kedde et al. 2010; Miles et al. 2012; Naudin et al. 2017; Goldstrohm, Hall, and McKenney 2018). PUM1 and PUM2 carry 76% sequence identity, bind to the same PRE sequence, and both are broadly and coincidentally expressed in tissues and cell lines (Spassov and Jurecic 2002; Goldstrohm, Hall, and McKenney 2018). Our PAC-Seq data emphasize that they coregulate the majority of target mRNAs. There exists a small subset of targets, however, which is impacted by only one PUM, similar to the effects observed by others (Yamada et al. 2020). The determinants that make those mRNAs particularly responsive to one PUM but not the other remain to be identified, but could include modulation by additional cis-acting sequence elements and/or trans-acting factors.

### Conservation of repression mechanisms among PUF proteins

Accumulating evidence indicates that the repression mechanism of PUF proteins is conserved. In addition to the data reported here, evidence from worms *(Suh et al. 2009)*, fruit flies *(Kadyrova et al. 2007; Weidmann et al. 2014; Arvola et al. 2020)*, fission yeast (Webster, Stowell, and Passmore 2019), and budding yeast (Goldstrohm et al. 2006) support the central role of CNOT in PUF-mediated repression. The interaction of the highly conserved RBD of PUF proteins with CNOT appears to be universal (Goldstrohm et al. 2006; Hook et al. 2007; Kadyrova et al. 2007; Suh et al. 2009; D. Lee et al. 2010; Weidmann et al. 2014). In contrast, the N-terminal repression domains that bind to CNOT, including RD3, are found in PUF proteins from organisms ranging from insects to vertebrates, but no homologous domains were detected in lower eukaryotes (Weidmann and Goldstrohm 2012; Goldstrohm, Hall, and McKenney 2018). In *Drosophila,* the sole Pumilio protein has three distinct N-terminal repression domains that can recruit CNOT to elicit repression (Weidmann and Goldstrohm 2012; Arvola et al. 2020), whilst an unrelated, unique region of the *S. pombe* PUF3 protein also binds CNOT (Webster, Stowell, and Passmore 2019). In addition to deadenylation, the involvement of the decapping factors in PUF-mediated repression is also conserved in humans (this study), *Drosophila (Arvola et al. 2020)*, and *S. cerevisiae* (Olivas and Parker 2000; Goldstrohm et al. 2006; Blewett and Goldstrohm 2012). Collectively, this research highlights the deep evolutionary conservation of the PUF regulatory mechanisms that dynamically control transcriptomes.

In summary, our results reveal the molecular mechanism by which PUM1&2 recruit the mRNA decay machinery to regulate gene expression. This new fundamental knowledge is anticipated to promote understanding of the crucial regulatory roles of PUMs in development and differentiation in diverse contexts including embryos and germline, hematopoietic, and nervous systems.

## Materials and Methods

### Plasmids and cloning

All oligonucleotides and plasmids used in this study are listed in **Tables S4 and S5**, respectively. The renilla luciferase (Rluc) altered specificity reporter, Rluc 3xPRE UGG, was created in psiCheck1 (Promega) as previously described (Van Etten et al. 2012). Nano-luciferase (Nluc) reporters were cloned by replacing Rluc coding sequence in psiCheck1 with the NanolucP coding sequence (Promega) using XbaI and SalI sites to generate plasmid pNLP. The Nluc-based reporters were then generated by inserting 3 wild type PREs (Nluc 3xPRE) or 3 mutant PREs (Nluc 3xPRE mt) into the XhoI and NotI sites of the 3′UTR using oligo cloning and inverse PCR (Van Etten et al. 2012; Bohn et al. 2018). The tethered function reporters Nluc 4xMS2 pA and Nluc 4xMS2 MALAT1, were similarly derived from pNLP as previously described (Abshire et al. 2018). The firefly luciferase (Fluc) plasmid, pGL4.13 (Promega), was used as a co-transfected control.

Expression plasmids for altered specificity (R6as) PUM1 and PUM2 full length and RBD constructs were previously described (Van Etten et al. 2012; Weidmann et al. 2014). PUM1 constructs with disease mutations (T1035S, R1139W, or R1147W (Gennarino et al. 2018)) were introduced into full length and RBD PUM1 R6as constructs using site-directed mutagenesis with oligos IE133/134, IE135/136, and IE137/138, respectively. The putative cap binding motifs (PUM1 W466G and PUM2 W350G) (Cao, Padmanabhan, and Richter 2010) were introduced in PUM1 and PUM2 full length R6as constructs using site-directed mutagenesis and oligos IE43/44 and IE120/121.

The tethered effector constructs PUM1 N (aa 1-827), PUM1 RD1 (aa 1-149), PUM1 RD2 (aa 309-459), PUM1 RD3 (aa 589-827), PUM1 RBD (aa 828-1175), PUM2 N (aa 1-704), PUM2 RD2 (aa 186-344), PUM2 RD3 (aa 471-704), PUM2 RBD (aa 705-1049) were Flexi cloned into mammalian expression vector pF5K (Promega) that contained the MS2 coat protein RNA-binding domain and a V5 epitope tag at the N-terminus. The tethered effectors pFN21A HT-MS2 and pFN21A HT-MS2 CNOT7 (Abshire et al. 2018) served as negative and positive controls, respectively, and were modified by inverse PCR with oligos IE177/178 and IE179/180, respectively, to include a V5 epitope tag.

The dominant negative CNOT7 (D40A, E42A) and CNOT8 (D40A, E42A) mutants were previously described (Piao et al. 2010; Van Etten et al. 2012). DCP2 (E148Q) dominant negative mutant (Chang et al. 2014) was generated by site directed mutagenesis using oligos IE128 and IE129 with template pcDNA3 myc DCP2 (provided by Dr. Jens Lykke Andersen, University of California, San Diego). For co-immunoprecipitation assays, pF5A vector (Promega) served as a negative control and plasmid pFN21A CNOT8 with an N-terminal V5 epitope tag was used as a positive control.

For the biochemical protein interaction assays, PUM constructs were created with N-terminal Maltose Binding Protein (MBP) tag and a cleavage site for human rhinovirus (HRV3C) protease, along with a C-terminal StrepII affinity tag. Human PUM constructs utilized in pull down assays included PUM1 RD1 (aa 1-149), PUM1 RD2 (aa 309-459), PUM1 RD3 (aa 589-827), PUM1 RBD (aa 828-1175), PUM2 RD2 (aa 186-344), PUM2 RD3 (aa 471-704), and PUM2 RBD (aa 705-1049). These inserts were cloned using Gibson assembly into the pnYC-pM vector that was linearized with NdeI (Gibson et al. 2009; Diebold et al. 2011; Arvola et al. 2020).

### Cell Culture and Transfections

Human HCT116 cells (ATCC) were cultured at 37°C under 5% CO2 in McCoy’s 5A media (Thermo Fisher Scientific) with 1x penicillin/streptomycin and 10% FBS (Invitrogen). Transfections were conducted using Fugene HD (Promega) according to the manufacturer’s protocol with a ratio of 4 μl of Fugene HD to 1 μg of plasmid DNA. For 96-well reporter assays, 5000 cells were plated in white-walled 96 well plates. 24 hr post seeding, the cells were transfected with 5 ng Fluc reporter plasmid and either 10 ng of Nluc or Rluc luciferase reporter plasmids (as indicated in the figures) and 85 ng of effector or control plasmid. For reporter assays involving overexpression of effectors and dominant negative mutants, cells were transfected with 5 ng Fluc reporter, 10 ng of Nluc reporter, 10 ng of effector and 37.5 ng of each of CNOT7 and CNOT8 mutant or 75 ng of the indicated negative control plasmid. For reporter assays involving overexpression of effectors and dominant negative DCP2 mutant, cells were transfected with 5 ng Fluc reporter, 10 ng of regulated Nluc reporter, 10 ng of effector and 75 ng of the DCP2 mutant plasmid or negative control plasmid, as indicated in the the respective figures. For reporter assays with effector plasmid titrations, cells were transfected with 5 ng Fluc reporter, 10 ng of Nluc reporter, and the indicated amount of effector plasmid. Total mass of transfected plasmid was balanced in each sample by addition of pF5A vector to a maximum total of 85 ng. For reporter assays that were conducted in six-well format, specifically those that included RNA analysis, each well was seeded with 2 × 10^5^ cells. After 24 hr, cells were transfected using 0.5 μg FLuc and 1.25 μg of Nluc reporter along with 1.25 μg of the effector or control plasmid indicated in the figure. For co-immunoprecipitation assays, 7×10^6^ HCT116 cells were seeded in a 100 mm dish and, after 24 hr, were transfected with 17 μg of the indicated plasmid DNA using Fugene HD.

### Reporter Gene Assays

The approach for luciferase-based PUM reporter gene assays was previously described (Van Etten et al. 2012; Van Etten, Schagat, and Goldstrohm 2013; Bohn et al. 2018). In all reporter assays, an internal control Fluc reporter is co-transfected with the indicated Nluc reporter so as to normalize for potential variation in transfection efficiency. Three types of luciferase-based reporter genes were used in this study: 1) PRE-based reporters that measure repression by endogenous PUMs; 2) Altered specificity PRE-based reporters that measure repression by transfected PUM effector proteins with programmed RNA binding specificity; 3) Tethered function reporters that measure the regulatory activity of transfected MS2-fusion effector proteins.

To measure repression by endogenous PUM1&2, a Nluc reporter mRNA bearing three wild type PREs in the 3′UTR (Nluc 3xPRE) was compared to a negative control reporter wherein the critical 5′-UGU sequence of each PRE was mutated to 5′-ACA (Nluc 3xPRE mt), as previously described (Van Etten et al. 2012; Bohn et al. 2018). This 5′-ACA mutation prevents binding by PUMs and eliminates repression activity (Van Etten et al. 2012; Bohn et al. 2018). Nluc and Fluc expression levels were measured using the Nano-glo Dual-luciferase reporter assay system (Promega) with a Glomax Discover luminometer (Promega).

To measure activity of wild type and mutant PUM1 or PUM2 effectors without interference from endogenous PUMs, an altered specificity (as) assay was previously developed (Van Etten et al. 2012; Weidmann et al. 2014). PUM1&2 were programmed to specifically bind a mutant PRE that has the 5′-UGU sequence changed to 5′-UGG (Van Etten et al. 2012). To do so, the RNA-recognition motif of the 6th repeat (R6) of each Pum-HD was changed to create PUM1 R6as and PUM2 R6as, which specifically bind and repress reporter mRNA bearing the PRE UGG (Rluc 3xPRE UGG) (Van Etten et al. 2012; Weidmann et al. 2014). This PRE UGG sequence is not recognized by the wild type endogenous PUMs (X. Wang et al. 2002; Van Etten et al. 2012). Rluc and Fluc activity was measured using the Dual-Glo Assay system (Promega). Expression of Halotag (HT)(Promega) served as a negative control in these experiments.

A tethered function approach (Jeffery Coller and Wickens 2002; Jeff Coller and Wickens 2007; Clement and Lykke-Andersen 2008) was used to measure the repressive activity of the N-terminus of PUM1 and PUM2, utilizing the Nluc 4xMS2 pA or Nluc 4xMS2 MALAT1 reporters that contained 4 stem-loop binding sites for the MS2 coat protein in their 3′UTR as previously described (Abshire et al. 2018). MS2 fusion protein effector constructs were expressed in conjunction with these reporters to measure their effect on gene expression. Nano-glo Dual-luciferase reporter assay was performed to measure the effects of each effector relative to the negative control MS2-Halotag (HT) fusion protein. As a positive control, experiments included the effector MS2-CNOT7 deadenylase, which causes robust repression and mRNA decay (Abshire et al. 2018).

Reporter assays were performed in 96-well format, unless noted otherwise. For reporter assays that included measurements of both reporter protein activity and mRNA level, a six-well format was used. Cells were trypsinized, counted, and re-seeded with the same number of cells per well (2 – 8 × 10^4^) into white 96-well plate prior to measurements using the dual luciferase assays.

Reporter data analysis was performed as previously described (Van Etten et al. 2012; Van Etten, Schagat, and Goldstrohm 2013; Bohn et al. 2018). For each sample, the relative response ratio (RRR) was calculated by normalizing NLuc, or Rluc, activity (measured in relative light units, RLU) to the corresponding Fluc value. These RRR values were then used to calculate fold change values between the indicated test conditions. For reporter assays measuring the activity of endogenous PUMs via PREs, fold change values were calculated from the RRR values for the PUM-regulated Nluc 3xPRE reporter relative to the unregulated mutant PRE reporter, Nluc 3xPRE mt. For RNAi experiments that tested the role of putative co-repressors, the effect of each RNAi condition on PRE/PUM-mediated repression was measured within that same RNAi condition, as previously established (Arvola et al. 2020). In this manner, the specific effect of co-repressor depletion on PRE/PUM activity is determined. The non-targeting control siRNA (NTC) served as negative control, whereas siRNAs for PUM1&2 served as a positive control. For altered specificity or tethered function assays, the fold change induced by an effector was calculated from RRR values relative to those for the negative control effector HT and MS2-HT respectively.

All reporter assays were performed in three independent experiments with a total of nine replicates. The resulting data are reported as mean log2 fold change values along with standard error of the mean (SEM). Statistical significance of comparisons, indicated in the figures and legends, was calculated using Student’s t test (paired, two-tailed), and the resulting p-values, number and type of replicates, and data are reported in the figures and **Table S1**.

### RNA interference

RNAi mediated knockdown experiments were performed using Dharmacon On-Target Plus Smartpool siRNAs (**Table S4**), which are optimized by the manufacturer to minimize potential off target effects. HCT116 cells were seeded at 2 × 10^5^ per well of a six-well plate. After 24 hr, the cells were transfected with 25 nM final concentration of the indicated siRNA using Dharmafect 4 (Dharmacon). A second transfection of siRNA was done after 24 hr for the assays involving the following knockdown conditions; CNOT7 and CNOT8 (combined knockdown), CNOT6 and 6L (combined knockdown), CNOT10 and CNOT11. 24 hr after the final siRNA treatment, cells were transfected with reporters as described above. Cells were harvested for assays 48 hr after the reporter transfection. Efficiency of each RNAi treatment was confirmed by western blot and/or RT-qPCR (described below).

### RNA Purifications and cDNA Preparation

For RNAi knockdown verification, RNA was purified from 2 × 10^6^ HCT116 cells harvested 48 hr following reporter transfection using the SimplyRNA cells kit and Maxwell RSC instrument (Promega) following the manufacturer’s protocol. Twice the amount of DNase was added for the on-bead DNase treatment. Purified RNA was eluted in 40 μl of nuclease-free water and then was quantified using a Nanodrop spectrophotometer (Thermo Fisher Scientific). For analysis of tethered function MS2 reporter mRNAs by RT-qPCR, RNA was purified from 2 × 10^6^ HCT116 cells 48 hr after reporter transfection. To ensure removal of potential DNA contamination, 3 μg of purified RNA was incubated with 3 units of RQ1 DNase (Promega) at 37°C for 30 mins. Next, the RNA was purified using a RNA Clean and Concentrator-25 spin-column (Zymo), eluted in 25 μl of water, and again quantitated. Reverse transcription was then performed as previously described using GoScript reverse transcriptase (Promega) with random hexamer primers (Van Etten et al. 2012; Arvola et al. 2020).

### Quantitative PCR

Quantitative PCR parameters are reported according to the MIQE guidelines in **File S1**, including primer set sequences, amplicon size, and amplification efficiencies (Bustin et al. 2009). To confirm RNAi depletion of CNOT6, CNOT6L, CNOT7 and CNOT8, RT-qPCR was performed using GoTaq qPCR Master mix (Promega). Cycling conditions were as follows: (i) 95 °C for 3 min, (ii) 95 °C for 10 s, (iii) 65 °C for 30s (for CNOT7 and CNOT8), 60 °C for 30s for (CNOT6 and 6L), or 65.5 °C for 30s (for internal control 18S rRNA) and (iv) 72 °C for 40 s. Steps ii–iv were repeated a total of 40 cycles. Melt curves were then generated using 65°C – 95°C as a range. For all primer sets, negative control reactions were performed in the absence of reverse transcriptase. Quantification cycles (Cq) were measured using the CFX Manager software (BioRad). Three experimental replicates were performed for each analysis. Fold change knockdown for each RNAi target gene was calculated relative to the NTC condition using the ΔΔCq method as previously described (Pfaffl 2001). To measure RNA levels of the NLuc 4xMS2 BS pA reporter, the following cycling conditions were used: (i) 95 °C for 3 min, (ii) 95 °C for 10 s, (iii) 63 °C for 30s and (iv) 72 °C for 40 s. Steps ii–iv were repeated a total of 40 cycles. Melt curves were generated as previously stated. The non-coding, non-adenylated 18S rRNA was chosen as the internal control for these experiments because of the potential for deadenylase knockdown to affect the levels of potential control mRNAs.

### Northern blotting

For Northern blot experiments, cells were transfected in six-well format with the indicated reporter gene, Nluc 4xMS2 pA. Purified total RNA (3 μg) was separated by formaldehyde-agarose gel electrophoresis with 1x MOPS buffer and then transferred to Immobilon NY+ member (Millipore) as previously described (Arvola et al. 2020). The RNA integrity and loading of the RNA in each sample was assessed by staining with ethidium bromide. The blot was pre-hybridized for 1 hr at 68 °C in 10 ml of UltraHyb buffer (Invitrogen) and then was probed with an antisense RNA Nluc probe that was transcribed with α-^32^P UTP (Perkin Elmer) as described (Arvola et al. 2020). After overnight hybridization, the blots were washed twice in 25 ml of 2x SSC, 0.1% SDS and then twice in 0.1x SSC, 0.1% SDS at 68°C for 15 min. Blots were visualized by phosphorimaging with a Typhoon FLA phosphorimager (GE Life Sciences) and quantified using ImageQuant TL software (GE Life Sciences). The 18S rRNA was detected using a 5′ ^32^P-labeled antisense oligodeoxynucleotide probe (listed in **Table S4**), prepared as previously described (Arvola et al. 2020). Pre-hybridized was done in 10 ml UltraHyb buffer for 1 hr at 42 °C and then the probe was incubated with the blot overnight. Blots were then washed twice with 25 ml 2×SSC containing 0.5% SDS for 30 min each wash at 42°C.

### Immunoprecipitation Assays

For co-immunoprecipitation analysis, 7×10^6^ HCT116 cells were seeded in a 100 mm dish and, after 24 hr, they were transfected with plasmids that expressed V5-tagged PUM1 RD3, PUM2 RD3, or CNOT8. Cells transfected with pF5A served as a negative control. After 48 hr, cells were lysed by passing through a syringe four times in buffer A (50 mM Tris-HCl pH 7.5, 300 mM NaCl, 0.5% Triton ×100, and 1 mM EDTA) with 2x Complete protease inhibitor cocktail (Roche). The resulting cell lysate was cleared of cell debris by centrifugation at 4°C for 10 mins at 20,000 x g. The supernatant was then centrifuged through a 0.45-micron filter at 4,000 x g and the resulting cell extract was measured using the DC-Lowry assay (BioRad). One milligram of total cellular protein was then incubated with 30 μl bed volume of anti-V5 beads (Sigma), which were pre-equilibrated in buffer A, 4 μg of RNase A (Promega), and 40 units of RNase ONE (Promega) for 1.5 hr at 4°C with end-over-end rotation. Beads were washed four times with buffer A for 5 min per wash with end-over-end rotation. Protein complexes were eluted in with 1x SDS PAGE loading dye with heating at 42 °C for 10 mins. The eluated protein was collected and then analyzed by western blot. RNase digestion of cellular RNA was verified by purifying RNA from supernatant after immunoprecipitation using Reliaprep RNA purification kit (Promega) and then analyzing it by denaturing formaldehyde-agarose gel electrophoresis followed by imaging of the ethidium bromide stained gel.

### Western blotting

For western blots, HCT116 cells from one well of a 96-well plate were lysed in 50 μl radioimmunoprecipitation assay (RIPA) buffer (25mM Tris-HCl pH 7.6, 150 mM NaCl, 1% NP-40, 1% sodium deoxycholate, 0.1% SDS) supplemented with 2x Complete protease inhibitor cocktail (Roche) and lysed using a cell disruptor. Lysates were cleared by centrifugation at 20,000 x g for 10 mins. Protein was quantified using Lowry DC Assay (BioRad) with a bovine serum albumin (BSA) standard curve. Equal mass of each cell extract (20 μg) was separated on a 4-20% Mini-PROTEAN TGX gel (BioRad) and then transferred to Immobilon P membrane (Millipore) followed by western blot with the primary antibodies indicated in the figures and detection by enhanced chemiluminescence (Pierce, Millipore). Antibodies are listed in **Table S5**.

### PAC-Seq Library Preparation and Analysis

Isoform level changes in gene expression in response to RNAi of PUM1, PUM2, PUM1&2, or CNOT subunits were measured using Poly(A) Click Seq (PAC-Seq) as previously described (Routh et al. 2017; Elrod et al. 2019). First, 2×10^5^ HCT116 cells were seeded in six-well format and then were transfected with the indicated siRNAs (25 nM) using Dharmafect 4 (6 μl per sample) as specified by the manufacturer (Dharmacon). Three biological replicate knockdowns were performed for each RNAi condition. After 48 hr, the cells were harvested and cell extracts prepared for western blot and RNA was extracted as described above. Knockdown of each RNAi target was verified by western blotting. Using the purified RNA, PAC-seq libraries were prepared as previously described (Routh et al. 2017). One μg of total RNA was reverse transcribed with the partial P7 adapter (Illumina_4N_21T) and dNTPs with the addition of spiked-in azido-nucleotides (AzVTPs) at 5:1. The p5 adapter (IDT) was Click-ligated to the 5′end of the cDNA with CuAAC. The cDNA was then amplified for 16 cycles using the Universal primer and 3′ indexing primer, followed by purification on a 2% agarose gel with extraction of the 200-300 base pair amplicon. Barcoded libraries were then pooled and sequenced with single-end, 150 base-pair reads on a Nextseq 550 (Illumina).

PAC-seq data was analyzed with the DPAC pipeline v1.10 (Routh 2019) using the exon centric approach with the –P –M –C –A –B and –D options. Alignments were to the hg38 genome using the Gencode v32 annotation. Results were filtered such that genes or exons required a minimum 5 mean reads in each sample, a 1.3 fold change, and an adjusted p-value < 0.05 to be scored as significantly differentially expressed. Genes with more than one poly(A)-site (PAS) additionally required a percent distal PAS usage change of 20 percent to be considered a change in 3′UTR isoform. The resulting differential expression analysis and statistics are reported in **Table S2**. Tests for statistical significance of overlapping gene expression sets were performed using the R package *SuperExactTest* v1.0.7.1 (M. Wang, Zhao, and Zhang 2015) which uses a hypergeometric test for multi-set overlap. The gene lists and statistics for the comparative analyses are reported in **Table S3**. Gene ontology analysis was performed using DAVID v6.8 (Dennis et al. 2003) and top enriched GO terms are listed in **Table S3**.

### In vitro pull-down assays

*In vitro* pull-down assays were performed to detect protein inactions between domains of PUM1 and PUM2 and the reconstituted, purified CNOT complex, as previously described (Arvola et al. 2020). StrepII- and MBP-tagged human PUM1 and PUM2 RD and RBD constructs were expressed in *E.coli* BL21 (DE3) Star cells (Thermo Fisher Scientific). Cells were grown in LB overnight at 37°C. Cells were lysed (8 mM Na_2_HPO_4_, 137 mM NaCl, 2 mM KH_2_PO_4_, 2.7 mM KCl, 0.3% (v/v) Tween-20, pH 7.4) and cleared. Lysate was incubated for 1 hr with 30 μl (50% slurry) of StrepTactin sepharose resin (IBA). Beads were then washed three times with lysis buffer and once with binding buffer (50 mM Tris–HCl pH 7.5, 150 mM NaCl). 50 μg of purified human CNOT complex was added to beads and incubated for 1 hr. For the intact CNOT complex, the protein components were as follows: CNOT1 (amino acids 1– 2376), CNOT2 (aa 1–540), CNOT3 (aa 1–753), CNOT10 (aa 25–707), CNOT11 (aa 257–498), CNOT7 (aa 1–285), CNOT6 (aa 1–558), CNOT9 (aa 1–299) (Raisch et al. 2019). For the pull-down analysis of CNOT modules, the following purified components were used: catalytic module contained NOT1 (aa 1093-1317), CNOT6 (aa 1-563), CNOT7 (aa 1-285); the NOT10/11 module contained CNOT1 (aa 1-682), CNOT10 (aa 25-707), CNOT11 (aa 257-498); the CNOT9 module contained NOT1 (aa 1351-1588), CNOT9 (aa 19-285); NOT module contained CNOT1 (aa 1833-2361), CNOT2 (aa 344-540), and CNOT3 (aa 607-753). Post incubation beads were washed three times with the binding buffer and then bound proteins were eluted with the binding buffer containing 2.5 mM D-desthiobiotin. Proteins were analysed by Coomassie blue stained SDS PAGE.

## Materials and data availability

All unique/stable reagents generated in this study are available from the Lead Contact without restriction. The [datasets/code] generated during this study are available at Gene Expression Omnibus (GSE159510).

## Acknowledgments

The authors wish to thank members of the Goldstrohm, Valkov, and Wagner laboratories for helpful suggestions and feedback. This research was funded grants from the National Institute of General Medical Sciences, National Institutes of Health [R01GM105707 to A.C.G.] and American Cancer Society Research Scholar Grant [RSG-13–080-01-RMC to A.C.G.]; Max Planck Society (to E.V.); NIH, National Institute of General Medical Sciences [R01-GM134539 to E.J.W.]; NIH, National Cancer Institute [R03-CA223893-01 to P.J.]. Publication costs were paid by University of Minnesota institutional funds.

## Author Contributions

I.E.: lead author, conceptualization, methodology, investigation, formal analysis, visualization, writing - original draft, review, editing

N.E.: methodology, formal analysis, visualization, editing

J.B.: resources, investigation, editing

C.C.: resources, investigation

Y.L.: resources, investigation

P.J.: investigation

A.L.: investigation

E.V.: conceptualization, experimental design, visualization, writing - review, editing, funding acquisition

E.J.W: conceptualization, methodology, funding acquisition, editing draft

A.C.G.: conceptualization, experimental design, analysis, visualization, writing - original draft, review, editing, project administration, funding acquisition

